# Dynamic Community Composition Unravels Allosteric Communication in PDZ3

**DOI:** 10.1101/2020.12.30.424852

**Authors:** Tandac F. Guclu, Ali Rana Atilgan, Canan Atilgan

## Abstract

The third domain of PSD-95 (PDZ3) is a model for investigating allosteric communication in protein and ligand interactions. While motifs contributing to its binding specificity have been scrutinized, a conformational dynamical basis is yet to be established. Despite the miniscule structural changes due to point mutants, the observed significant binding affinity differences have previously been assessed with a focus on two α-helices located at the binding groove (α_2_) and the C-terminus (α_3_). Here, we employ a new computational approach to develop a generalized view on the molecular basis of PDZ3 binding selectivity and interaction communication for a set of point mutants of the protein (G330T, H372A, G330T-H372A) and its ligand (CRIPT named L_1_ and its T-2F variant L_2_) along with the wild type (WT). To analyze the dynamical aspects hidden in the conformations that are produced by molecular dynamics simulations, we utilize variations in community composition calculated based on the betweenness centrality measure from graph theory. We find that the highly charged N-terminus which is located far from the ligand has the propensity to share the same community with the ligand in the biologically functional complexes, indicating a distal segment might mediate the binding dynamics. N- and C-termini of PDZ3 share communities, and α_3_ acts as a hub for the whole protein by sustaining the communication with all structural segments, albeit being a trait not unique to the functional complexes. Moreover, α_2_ which lines the binding cavity frequently parts communities with the ligand and is not a controller of the binding but is rather a slave to the overall dynamics coordinated by the N-terminus. Thus, ligand binding fate in PDZ3 is traced to the population of community compositions extracted from dynamics despite the lack of significant conformational changes.

## INTRODUCTION

PDZ domains are abundant in various complexes, and the well-studied PSD-95 is from the MAGUK family with a PDZ-SH3-GK pattern. The third PDZ (PDZ3) of this complex is important for the formation of this supramodule and its binding to C-terminal ligands.^1–6^ Most of the PDZ domains have two α-helices and six β-strands. However, PDZ3 is unique with an extra α-helix (α_3_) located at its C-terminus. α_3_ has been shown to be important in the folding of PDZ3 as well as its interactions with SH3 and ligands.^1, 5, 7–11^

The structure of wild-type (WT) PDZ3 has been solved with its cognate ligand CRIPT;^12, 13^ it has been demonstrated that PDZ3 binds specific motifs of ligands.^14^ The ligands are classified by amino acid type at the second position from the C-terminus (−2 position), with those having Thr/Ser defined as Class-I; this class contains CRIPT.^13^ Class-II has hydrophobic, and Class-III has Asp/Glu amino acids at this position.^14^

In this study, we focus on the WT, G330T, H372A and G330T-H372A (double mutant, DM) variants in complex with the ligand CRIPT (L_1_) and its T-2F form (L_2_).^15–17^ These theoretical variants of PDZ3 have been investigated by using mutational scans, their functionality have been assessed by experimental binding constants, and their crystal structures have been deposited.^16–19^

In an elegant work on PDZ, Raman et al. have shown that the adaptive evolutionary pathway for switching ligand binding from Class I to Class II utilizes a class bridging mutation such as G330T while the H372A mutation may only be gained at a second step.^17^ Hence, the binding experiments display that WT forms a functional complex only with L_1_. On the other hand, the G330T single mutant prefers binding to both L_1_ and L_2_; however, H372A single mutant and DM proteins favor only L_2_-binding.^16^ Thus, of the eight possible complexes, WT_L1_, G330T_L1_, G330T_L2_, H372A_L2_ and DM_L2_ are functional, while WT_L2_, H372A_L1_ and DM_L1_ forms are unfavorable (the two respective groups are colored green and red in Table 1 and this color coded labelling has been maintained throughout this manuscript).

**Table 1.** The abbreviation and PDB IDs of the PDZ3 structures utilized for this study. Green and red colors indicate favorable and unfavorable complexes, respectively.

| Abbreviation | PDB ID | Structure Details |
| --- | --- | --- |
| WT<sub>L1</sub> | 5HEB | The third PDZ of PSD-95 in complex with CRIPT |
| WT<sub>L2</sub> | 5HED | The third PDZ of PSD-95 in complex with mutated CRIPT (T-2F) |
| G330T<sub>L1</sub> | 5HEY | The third PDZ of PSD-95 (G330T single mutant) in complex with CRIPT |
| G330T<sub>L2</sub> | 5HF1 | The third PDZ of PSD-95 (G330T single mutant) in complex with mutated CRIPT (T-2F) |
| H372A<sub>L1</sub> | 5HFB | The third PDZ of PSD-95 (H372A single mutant) in complex with CRIPT |
| H372A<sub>L2</sub> | 5HFC | The third PDZ of PSD-95 (H372A single mutant) in complex with mutated CRIPT (T-2F) |
| DM<sub>L1</sub> | 5HFE | The third PDZ of PSD-95 (G330T-H372A double mutant) in complex CRIPT |
| DM<sub>L2</sub> | 5HFF | The third PDZ of PSD-95 (G330T-H372A double mutant) in complex with mutated CRIPT (T-2F) |

To understand the molecular basis of binding, we investigate the previously studied structural segments, which are the N and C termini, α_2_ helix and the aforementioned α_3_ helix (Figure 1a).^5, 9–11, 20–22^ The dynamics of the highly charged N-terminus region (Figure 1a) has recently been shown to be important in the binding mechanism, especially due to its electrostatic contributions to the total free energy.^20^ The effect of the N-terminus might be partnered with that of the C-terminus.^21^ α_2_ helix (Figure 1a) lines up the ligand, and its effect has been investigated by deep mutational scan.^22^ Mutants of H372, residing on the α_2_ helix, have been shown to cause significant binding constant shifts when coupled with other point mutations, and this effect is conserved in other PDZ domains.^16, 22^ In PDZ3, H372 is in direct contact with the T-2 residue of L_1_ implying an essential role for ligand binding.^12, 16, 22^

**Figure 1.**
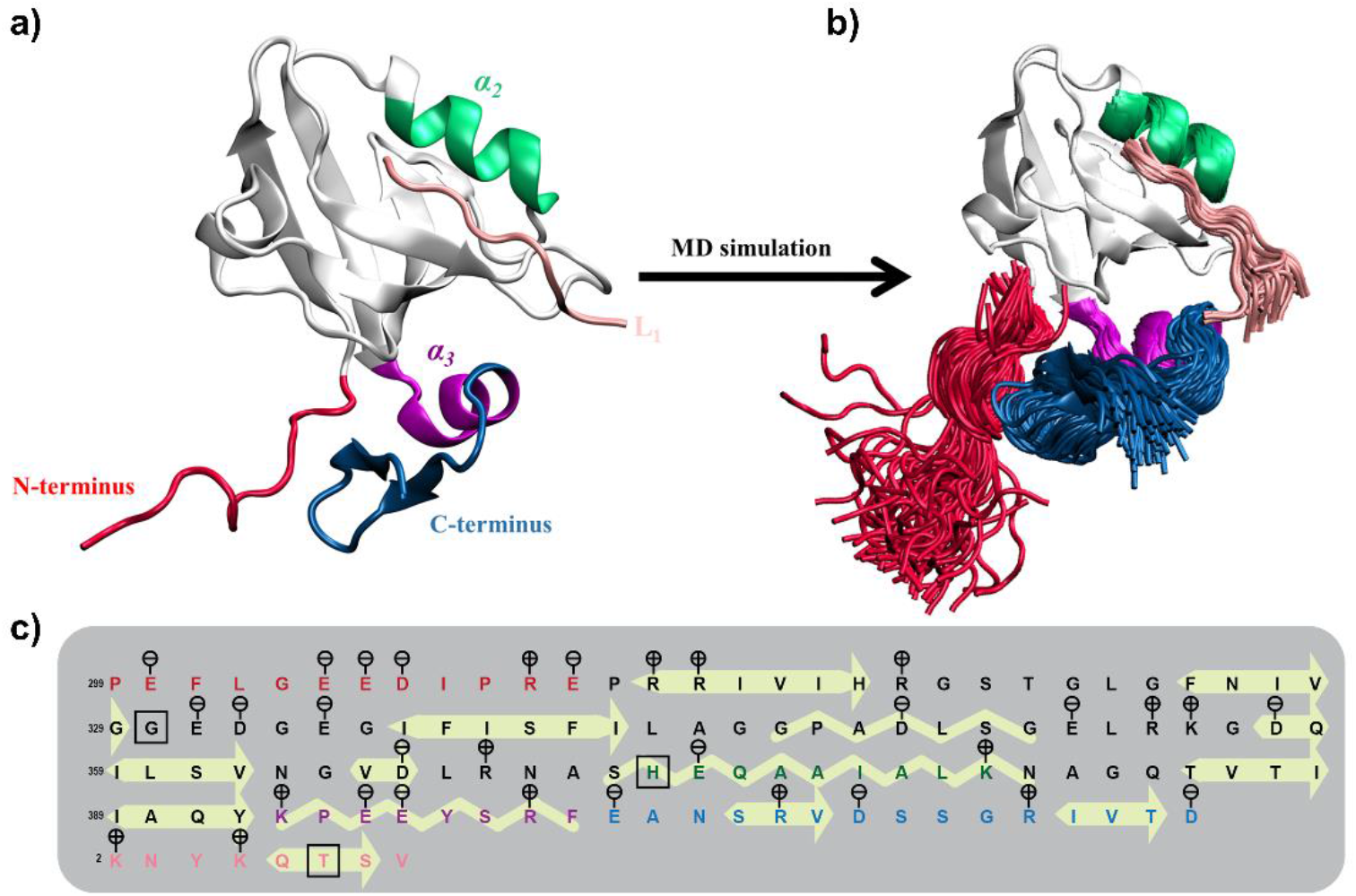
**a** Structure and sequence of WT_L1_ complex (PDB code 5HEB). N-terminus (residues 299-310), α_2_(residues 372-380), α_3_(residues 393-400), C-terminus (residues 401-415) and ligand (residues 2-9) are illustrated in red, green, purple, blue and magenta, respectively. **b** 120 snapshots from the MD simulations are illustrated for the highlighted structural elements. **c** PDZ sequence and details of the structural elements. Charges of residues are marked in circles above amino acid symbols. Yellow shapes indicate secondary structures, arrow for β-strands and zigzag for α-helices. Squared-residues display the mutation positions, which are G330, H372 in the protein and T-2 in the ligand.

In this work, we employ graph theory to investigate the communication between these functionally important structural segments.^23–26^ Coarse-graining the protein structure and projecting it to a residue network (RN) reduces that structure to nodes (vertices) and edges (links).^27–29^ In an RN, centrality of nodes reveals the residues that manipulate information flow, and identifying residues with high centrality provides a profile of biological function and evolutionarily conservation.^26, 30–33^ On the other hand, focusing on edge centrality in order to detect ‘communities’ has been put forth as an approach to distinguish modular units with interdependent functions for any network,^34–36^ and these ideas have been applied to residue networks to shed light on communication patterns between structural segments.^34–36^

Here, we analyze molecular dynamics (MD) simulations to gather a range of conformations sampled by PDZ3 with a novel approach (Figure 1b). We first apply network analysis on extracted MD snapshots;^37^ then, we track the dynamical changes of community members to their structural and functional origins. Community analysis reveals hidden allostery in protein structures by assessing the communication scenarios between the structural segments.^35, 36^ We propose that this method may be used to cluster the conformational dynamics of protein structures, and to infer information flow underlying functional mechanisms.

## MATERIALS AND METHODS

### System construction for molecular dynamics simulations

The MD simulations and their detailed energetic analyses have recently been reported elsewhere.^20^ Briefly, protein structures are downloaded from the protein data bank (PDB)^38^ (Table 1), and missing residues of ligand are inserted by using SWISS-MODEL^39^ server. Equal lengths of the structures are used whereby residue index ranges are 299-415 for the protein and 2-9 for the ligand (see Fig. 1c for the sequence). The structures are solvated in a rectangular water box with a minimum distance of 10 Å from all directions of the protein; KCl ions are added to the complex to achieve isotonic (150 mM) conditions. Simulations are carried with the NAMD^40, 41^ software, and CHARMM36^42^ force-field is utilized for parameters and topologies. The force-field parameter settings are as follows: Particle mesh Ewald (PME) grid size is 1 Å, cut-off distance is 12 Å, pair list distance is 13.5 Å, and switching is on with a distance of 10 Å. For all complexes, the minimization process is 10000 timesteps, then a 200-ns of simulation time is performed at 310 K constant temperature and under 1 atm constant pressure. Simulation of each complex is repeated to enhance sampling. The first 80 ns portion of all MD trajectories are discarded for equilibration, and the last 120 ns portions are used in all analyses; therefore, snapshots of 240 ns long MD simulation for WT, G330T, H372A and DM in complex with L_1_ and L_2_ are utilized for further analyses. Additionally, MD simulations for truncated (Δ) complexes are performed for 50 ns, and the whole trajectory is utilized for calculations in these systems with minimal amounts of fluctuations.

### Construction of Residue Networks

To construct a graph of the protein structure, C_β_ of each residue (C_α_ for glycine) is taken as a node to preserve side-chain sensitivity in calculations. Nodes within a 6.7 Å distance are taken as interacting, and an edge is assigned between them. The cut-off distance of 6.7 Å is chosen for linking the first coordination shell of C_β_ atoms in radial distribution function (RDF) which belongs to adjoint residues and other residues that locate close to the central residue;^25, 26^ for a detailed account of the choice of cutoffs in residue networks, see ref^43^. This cut-off is validated by the RDF analysis of our MD simulations, which shows that the dynamics and sampled system do not affect the overall coordination shell location (Figure S1). RNs are unweighted, undirected, and they do not have self-loops or parallel edges.

### Calculation of betweenness centrality

Capability of nodes or edges to manipulate information flow in a network is assessed by betweenness centrality (BC).^44, 45^ Assuming that information exchange happens through the shortest paths on a network;^25^ the BC of an arbitrary element (node or edge) is calculated by,

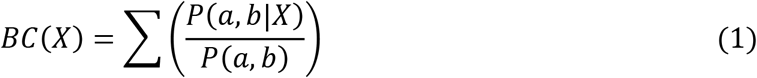

where *P(a, b|X*) is the number of the shortest paths travelled through this element, and *P(a, b)* is the number of all shortest path between the nodes *a* and *b*. BC is normalized by (*n* − 1)(*n* − 2)/2 for nodes and *n*(*n* − 1)/2 for edges, where *n* is the number of nodes in the network. The range of BC is [0, 1]; if all the shortest paths traverse through the certain element (*X*), it is 1. Hence, BC gives both local and global information about a graph, and it has been effectively used for structural and functional assessment of proteins.^26, 31–33, 35, 36^

### Detection of communities and structural origins of members

Girvan-Newman algorithm is employed to detect communities.^34, 46^ A community is a group of nodes that are connected to each other without having an edge to the nodes out of this group. Hence, members of same community are in cooperation, which, in the context of proteins, translates to function. The algorithm searches for communities by breaking the most central (highest BC) edge in each iteration, and the occurring communities (*Ω*) are utilized for further analyses. Along with the total edge count, number of removed edges until community separation is achieved proves informative about the state of the whole system. To scrutinize the structural origins of community members, we devise a simple algorithm. For an RN belonging to an MD snapshot, each community in *Ω* is checked whether at least one member from desired two different structural segments is located in the same community; if they are, the score for that *Ω* is 1, otherwise it is 0. The sum of scores is normalized by the total number of MD snapshots for average results. Accordingly, 240 snapshots from MD simulations (extracted from the equilibrated portions of the trajectories at 1 ns intervals) of each complex are utilized for the average results, and 24 snapshots (extracted at 10 ns intervals) are used for the detailed investigation of the conformations.

### Visualization of community dynamics on three-dimensional protein structure

To visualize the dynamical shifts in shared communities, we apply the following procedure: Our aim is to color each residue according to its persistence in a given community. Thus, we first select three residues that predominantly remain in separate communities and are in rigid structural elements. Here we have selected I316 in β_1_, A375 in α_2_ and F400 in β_3_. These residues are attributed the colors red (R), green (G) and blue (B), respectively. After performing community detection at each snapshot for a given *Ω*, we assign an attribute of R, G, B or null to each residue at every time point *t* in vector **c**_*i*_(*t*) such that if the residue is in the same community with I316, **c**_*i*_(*t*) = [1 0 0], if it is with A375, **c**_*i*_(*t*) = [0 1 0], if it is with F400, **c**_*i*_(*t*) = [0 0 1]; **c**_*i*_(*t*) = [0 0 0] otherwise. The color of the protein is accumulated in the color matrix **C** of dimensions *n* × 3 with each row holding the RGB color code of the residue, 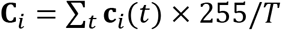, where *T* is the total number of time points so that the normalization by 255/*T* allows the use of decimal RGB code. The protein three-dimensional structure is then colored according to these values. As a result, residues always co-inhabiting a community with I316, A375 or F400 will have a pure R, G or B color, but those switching between regions will have blended colors, e.g., a residue spending half of its time in the same community with I316 and the other with A375 will appear yellow. Finally, those residues that are never clustered with any of the three will be colored black. In this study, this visualization has been applied to community sizes of *Ω* = 4.

## RESULTS AND DISCUSSION

In a recent study, we have investigated the relative binding energies of PDZ3 complexes using classical MD trajectory analyses and free energy perturbation calculations, and we have shown that the generated MD trajectories sample adequate conformational range to reproduce the experimentally measured binding affinities.^17, 20^ In particular, we have found that the interplay between protein-ligand hydrogen bonding, solvation free energy of the complex and the dynamics of the highly charged N-terminus are the three main factors that determine the binding fate of the ligand.^20^ In fact, it has been possible to compute the free energy difference values corroborating experimentally measured binding constants using various advanced simulation techniques and to break down the contribution of each of these factors by a simplified model. However, the energy values do not suffice to draw conclusions for a mechanistic interpretation of the findings. It is of interest to pinpoint mechanisms controlling these binding events by determining the role of key regions in this notoriously allosteric protein.

### Betweenness centrality unveils hinge residues affecting function of the complex

For this purpose, we first investigate if the BC of residues calculated as an average over the snapshots obtained through MD provide information additional to their mean-squared fluctuations (MSF). Selected case of H372A_L2_ is displayed in Figure 2a and data for all cases are provided in Figure S2. It is well known that high MSF residues are in mobile regions, mostly on solvent exposed loops, while residues in secondary structural elements have low MSF. Meanwhile, BC works at the resolution of single residues and reveals that even in flexible loops there are residues with high BC, undertaking hinge roles in PDZ binding mechanics (Figure 2a). We note that for the same mutant, binding different ligands may significantly shift the centrality of resides, best displayed by the ΔBC curves as exemplified for H372A in Figure 2b wherein the N-terminus residues G303 and E304 as well as the turn residue S409 have much increased centrality in the functional H372A_L2_ compared to the low binding affinity complex H372A_L1_(Figure 2b).

**Figure 2.**
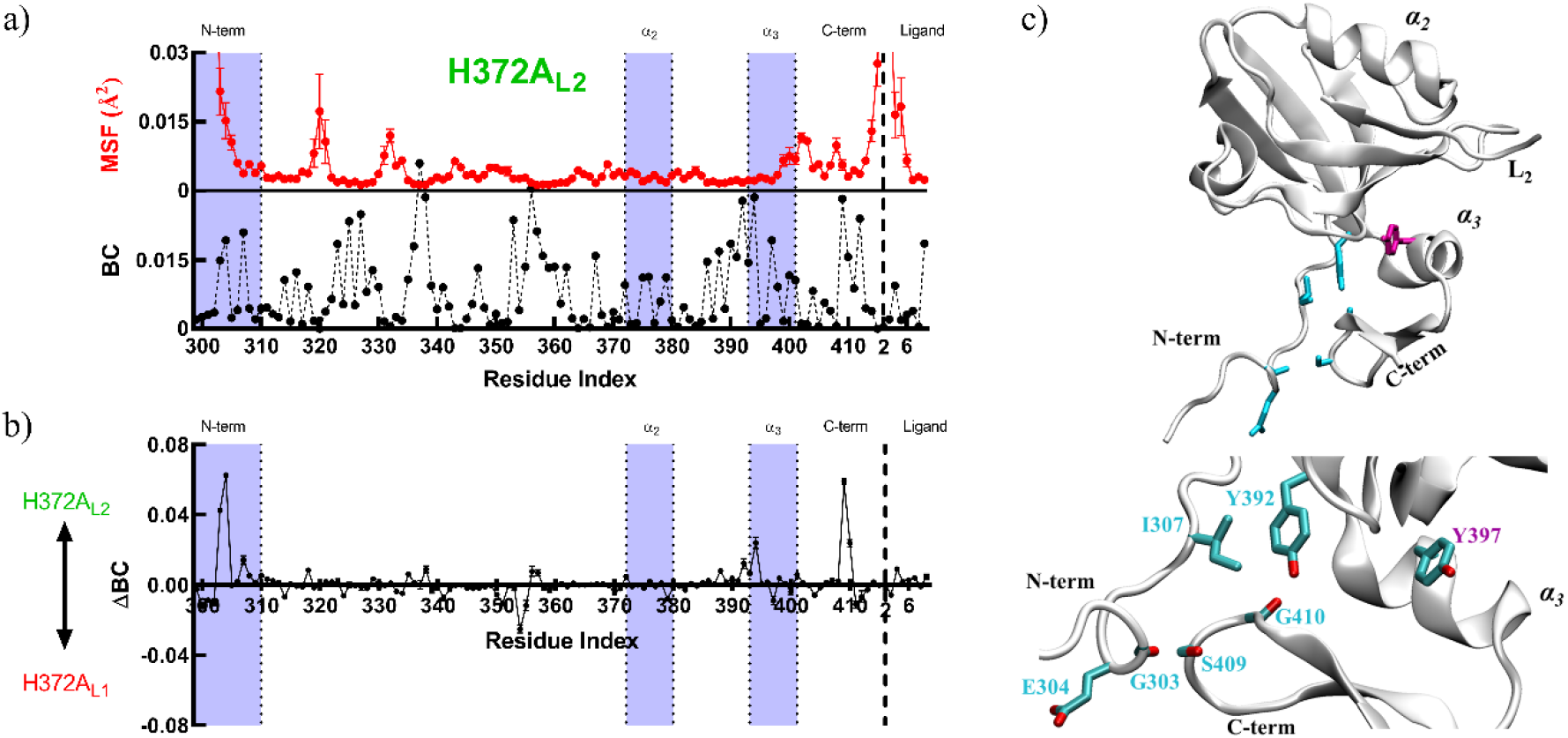
**a** Sample MSF and BC for H372A_L2_, full set shown in Figure S2. Structural segments are labeled at the top of the graph, and segments of interest are highlighted in purple. MSF results are averages over six chunks of 40 ns each obtained from the equilibrated duplicate MD trajectories. **b** ΔBC between L_1_ and L_2_ bound forms of H372A. Along with α_3_, N and C termini are more central in H372A_L2_. **c** Residues with |ΔBC| > 0.04 emerging in any of the studies complexes are mapped on the three-dimensional PDZ structure; residues with large changes in full length proteins are shown in cyan; Y397 (magenta) is highlighted only in truncation mutants along with Y392 and S409 that appear in both types of systems. All these residues reside on the N-terminus, C-terminus or the α_3_ helix; their locations and interactions shown in detail below.

In fact, these residues arise frequently amongst those with the largest ΔBC in all variants, whose locations and interactions are displayed in Figure 2c indicating the high centrality of N and C termini. We find G303/E304/I307 in the N-terminus, Y392 at the beginning of the α_3_ helix, and S409/G410 in the C-terminus to significantly shift their centrality depending on the variant. Moreover, in truncation simulation series, PDZ3^Δ^, whereby the N-terminus has been deleted, the BC of C-terminus region drops in all cases, indicating that N and C termini interaction is substantial for the formation of the PDZ complexes. In these truncation variants, Y392, Y397 and S409 commonly lose centrality. Note that Y397 resides in the middle of the α_3_ helix and does not directly interact with the N-terminus in full length proteins providing an example of how the lost interactions lead to a domino effect that reflect into the bulk of the protein in the communication. Finally, note how for all complexes, BC of Y392 shifts, emphasizing the function of this residue on information flow. Y392 is located at the beginning of α_3_, and it is central in all complexes; along with the high centrality of N/C termini, this residue appears to hold a mediator function for PDZ3 without conferring ligand specificity.

While residues with high BC changes in the different complexes may indicate regions to target to alter the function of a protein, they alone cannot pinpoint how function is dynamically orchestrated. For this purpose, we turn to analyze how communities of residues are structured.

### Number of broken edges and size of the communities illustrate diverse organizations of PDZ3

Rather than individual residues, we now focus on the edges of the PDZ3-RNs, i.e., the interactions between residue pairs. In particular, an analysis of groups of residues working together is deciphered by studying the composition of communities formed by structural segments during the course of the time in the trajectories. To detect the communities, the edges are removed one-by-one hierarchically, starting from the most central, until a group of residues separate out into a disconnected community. The total number of edges in the range of 360-380 show that RNs of PDZ3 are very sparse (Table 2), compared to the maximum number of possible edges, which is *n*(*n* − 1)/2 = 7750 where *n* = 125 is the number of nodes. Considering the sparse character of the networks and the prior assessment of the communities, a community window of *Ω* = 3-6 is selected for detailed study. At *Ω* = 2, the flexible N-terminus and protein body are grouped separately as a trivial result; for *Ω* > 6 single residue communities start to dominate.

**Table 2.** Averages for total number of edges, and number of broken edges to achieve a community of size *Ω.*

| System label | Total number of edges | Average number of broken edges |  |  |  |
| --- | --- | --- | --- | --- | --- |
| | | $\Omega = 3$ | $\Omega = 4$ | $\Omega = 5$ | $\Omega = 6$ |
| **WT<sub>L1</sub>** | $365 \pm 1$ | $31 \pm 1$ | $51 \pm 1$ | $63 \pm 1$ | $72 \pm 1$ |
| **WT<sub>L2</sub>** | $364 \pm 1$ | $29 \pm 1$ | $50 \pm 1$ | $63 \pm 1$ | $73 \pm 1$ |
| **G330T<sub>L1</sub>** | $363 \pm 1$ | $26 \pm 1$ | $47 \pm 1$ | $60 \pm 1$ | $70 \pm 1$ |
| **G330T<sub>L2</sub>** | $364 \pm 1$ | $30 \pm 1$ | $49 \pm 1$ | $62 \pm 1$ | $71 \pm 1$ |
| **H372A<sub>L1</sub>** | $366 \pm 1$ | $36 \pm 1$ | $54 \pm 1$ | $64 \pm 1$ | $71 \pm 1$ |
| **H372A<sub>L2</sub>** | $367 \pm 1$ | $27 \pm 1$ | $48 \pm 1$ | $60 \pm 1$ | $71 \pm 1$ |
| **DM<sub>L1</sub>** | $376 \pm 1$ | $38 \pm 1$ | $57 \pm 1$ | $68 \pm 1$ | $77 \pm 1$ |
| **DM<sub>L2</sub>** | $374 \pm 1$ | $40 \pm 1$ | $56 \pm 1$ | $67 \pm 1$ | $75 \pm 1$ |

To provide an insight on how the community sharing and BC data lead to complementary information, in Figure 3 we display as heatmaps the BC value at each instant of the trajectories, accompanied below them with the fraction of time the N-terminus and the ligand share a community for the range of *Ω* = 3-6. The ligand has low BC in all cases (black stripes in the topmost part of the figures), and several residues of the N-terminus has high BC (light colored instants in the lowest part of the figures). Nevertheless, these two regions frequently share a community for *Ω* = 3, but their communities may further separate out depending on the system studied for larger *Ω* values (checkerboxes in Figure 3).

**Figure 3.**
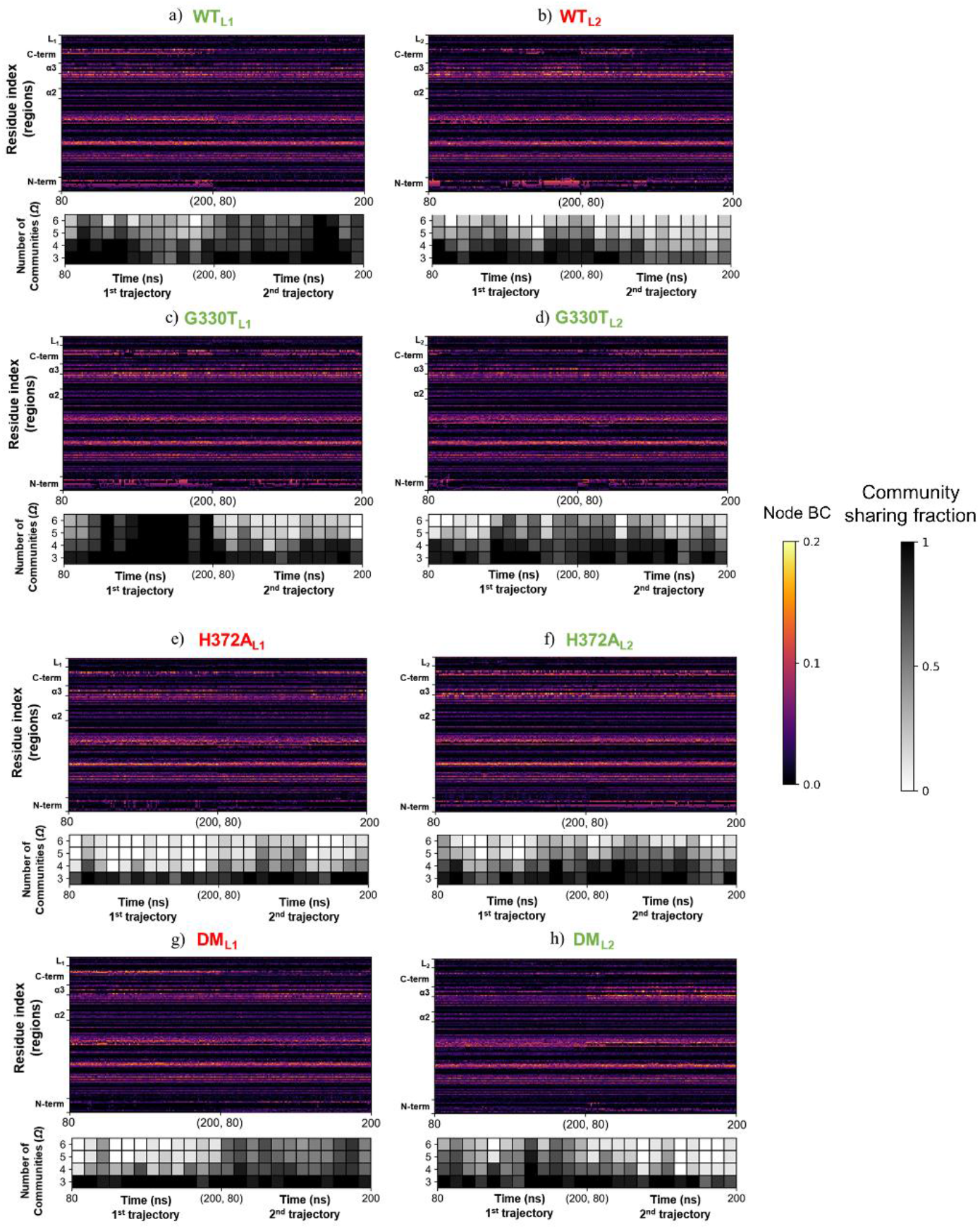
Heatmap of Node BC (colored; labelled by structural segments instead of residue indices) paired to the fraction of community sharing of N-terminus and ligand for *Ω* = 3-6 (grayscale cells, each cell average of 10 snapshots) throughout the MD trajectories. The results for two replica trajectories are concatenated, with time points covering 80-200 ns for each as shown in x-axis labels.

The total number of edges averaged over the trajectory snapshots as well as the number of edges needed to be broken to reach a given community size are listed in Table 2. They are ~365 in WT and the single mutants, while DM forms have ~375 edges. This is a result of the close-knit character of DM complexes, having more hydrogen bonds at the binding pocket and low overall solvent accessibility, as quantified in detail in our previous study.^20^ Although the averages are similar, the number of broken edges fluctuate over the course of the trajectories (Figure S3). These changes indicate that the configurations for information flow are modified throughout the MD trajectories. When we focus on the size of the communities, we find that at *Ω* = 3 there is one large community having 60-100 residues accompanied by two smaller ones (compare the green curves to red and blue curves in Figure S4 for *Ω* = 3). As more edges are removed, the variations at different time points smooth out and the size of the communities gets more similar, with the largest community having 20-60 members and the smallest one having a few members (see Figure S3 for *Ω* = 6).

The composition of the communities is investigated by assessing the structural origins of its members. The fraction of being a member of the same community is calculated for all available pairs between five structural segments that have been determined in the BC analysis to be essential for PDZ3; namely the N/C termini, α_2_/α_3_ helices and ligand (refer to Figure 1 for residue indices). Note that two segments are classified to be members of the same community if at least one residue is shared; therefore, the fractions do not sum to 1. The results for all pairs of structural segments and community sharing for the range of *Ω* = 3-6 are listed in Table S1 and that for the particular case of the N-terminus and ligand is in Table 3.

**Table 3.** Fraction of instances that N-terminus and ligand co-inhabit a community.*

| System label | $\Omega = 3$ | $\Omega = 4$ | $\Omega = 5$ | $\Omega = 6$ |
| --- | --- | --- | --- | --- |
| **WT<sub>L1</sub>** | **$0.93 \pm 0.02$** | **$0.84 \pm 0.02$** | $0.66 \pm 0.03$ | $0.56 \pm 0.03$ |
| **WT<sub>L2</sub>** | **$0.84 \pm 0.02$** | $0.63 \pm 0.03$ | $0.32 \pm 0.03$ | $0.14 \pm 0.02$ |
| **G330T<sub>L1</sub>** | **$0.90 \pm 0.02$** | **$0.80 \pm 0.03$** | $0.44 \pm 0.03$ | $0.26 \pm 0.03$ |
| **G330T<sub>L2</sub>** | **$0.93 \pm 0.02$** | **$0.79 \pm 0.03$** | $0.55 \pm 0.03$ | $0.48 \pm 0.03$ |
| **H372A<sub>L1</sub>** | **$0.88 \pm 0.02$** | $0.31 \pm 0.03$ | $0.14 \pm 0.02$ | $0.09 \pm 0.02$ |
| **H372A<sub>L2</sub>** | **$0.88 \pm 0.02$** | **$0.70 \pm 0.03$** | $0.35 \pm 0.03$ | $0.15 \pm 0.02$ |
| **DM<sub>L1</sub>** | **$0.97 \pm 0.01$** | $0.56 \pm 0.03$ | $0.36 \pm 0.03$ | $0.33 \pm 0.03$ |
| **DM<sub>L2</sub>** | **$0.90 \pm 0.02$** | $0.50 \pm 0.03$ | $0.32 \pm 0.03$ | $0.21 \pm 0.03$ |

\*values greater than 0.7 shown in bold; those less than 0.5 are colored blue.

First and foremost, N-terminus – C-terminus – α_3_ are located in the same community even for the largest value considered of *Ω* = 6 with fraction of time they spend together in pairs exceeding values 0.7 throughout and nearly equal to 1 in most cases (Table S1). In addition to the previously mentioned node BC results, the high fraction of N/C termini co-inhabiting the same community along with α_3_ emphasizes the interaction between the two segments is critical for all complexes. The two termini exert a clamping effect that facilitates the overall dynamics.^21^ Considered with the high node BC (Figure 2, Figure S2), α_3_ acts as a hub between structural segments, jointly with its neighboring Y392. Similarly, α_2_ that hosts H372 and the ligand reside in the same community throughout the trajectories (Table S1), an expected result since α_2_ lines the ligand (Figure 1). A series of manuscripts discuss the allosteric communication between the ligand binding site and the α_3_ helix.^5, 7–11^ Our analyses show that the two regions share a community with a fraction higher than ~0.6 in all PDZ complexes even at *Ω* = 6. However, community sharing between α_3_ and α_2_ is relatively low, particularly for WT_L1_ and G330T single mutation systems.

The fraction of community sharing between N-terminus and ligand is a measure of the extent to which the distal mobile region affects ligand binding (Table 3). We find that this couple tends to be in the same community in favorable complexes up to *Ω* = 5, except for DM_L2_. Also, the analyses between N/C termini, α_2_ and α_3_ do not indicate this specificity, which means that N-terminus communicates directly with the ligand. The effect is due to the long-range electrostatic interactions between the N-terminus having −4 net charge and the ligand with +2 charge. It is filtered through the protein core, and its range is manipulated by point mutations at positions 330 and 372. As a result, the flexible N-terminus interacts with the ligand and affects binding specificity, significantly for WT and single mutations. We have shown previously^20^ that the strong ligand binding preference of DM is dominated by the tight structure attained by this variant upon ligand binding that affects the overall solvation free energy change due to the reduction in solvent accessible surface area and additional hydrogen bonds counts at the binding site. The preference of L_2_ binding over L_1_ in this case is explained by the enhanced BC of the α_3_ hub that manifests itself as the tendency of the N-terminus, C-terminus and α_3_ to coinhabit the same community even at *Ω* = 6 (fraction of finding all three together is 0.88 for DM_L2_ vs. only 0.47 for DM_L1_.)

The analyses of the remaining segments do not provide information on binding specificity of PDZ3. For example, the C-terminus does not communicate with structural segments other than the N-terminus. Although, PDZ3-specific α_3_ has a higher fraction of communication with ligand and α_2_, the values do not differentiate functional complexes implying that α_3_ does not directly affect ligand selectivity. We therefore find that the N-terminus is the main region affected by the single residue changes that lead to ligand selectivity in PDZ3.

In fact, removal of the N-terminus significantly alters the community structure of PDZ3. In Table S2 we display the community sharing fractions for all pairs of the remaining regions in PDZ3^Δ^ simulation series where the values that differ from the full length PDZ3 variant (Table S1) by more than the error margins are colored in red. We find that the already low community sharing between C-terminus and α_2_/ligand in the full length PDZ3 is nearly completely lost even for *Ω* = 3 in all variants except H372A_L2_ where there is slight increase in communication between these regions. Moreover, focusing on the central role attributed to α_3_ to ligand binding (Figure 4), their community structure is also significantly altered, especially for WT_L2_ whose communication with the ligand is substantially disrupted for WT_L2_, but reinforced for H372A_L1_, and H372A_L2_. The N-terminus is almost always grouped together with α_3_ and ligand for *Ω* = 3 but is the first region to separate out for *Ω* > 3 (Figure 4, empty squares). If this region were not to affect the community dynamics of α_3_ and ligand, one would expect α_3_ and ligand to have the same occupancy fractions in PDZ3 and PDZ3^Δ^ simulations (Figure 4 circles and triangles). Indeed, we have discussed how the binding affinity of the DM variants is mainly governed by the tight overall structure, unlike the rest of the systems. Therefore, as expected, the absence of the N-terminus has no effect on the community structure of α_3_ and ligand in the DM systems.

**Figure 4.**
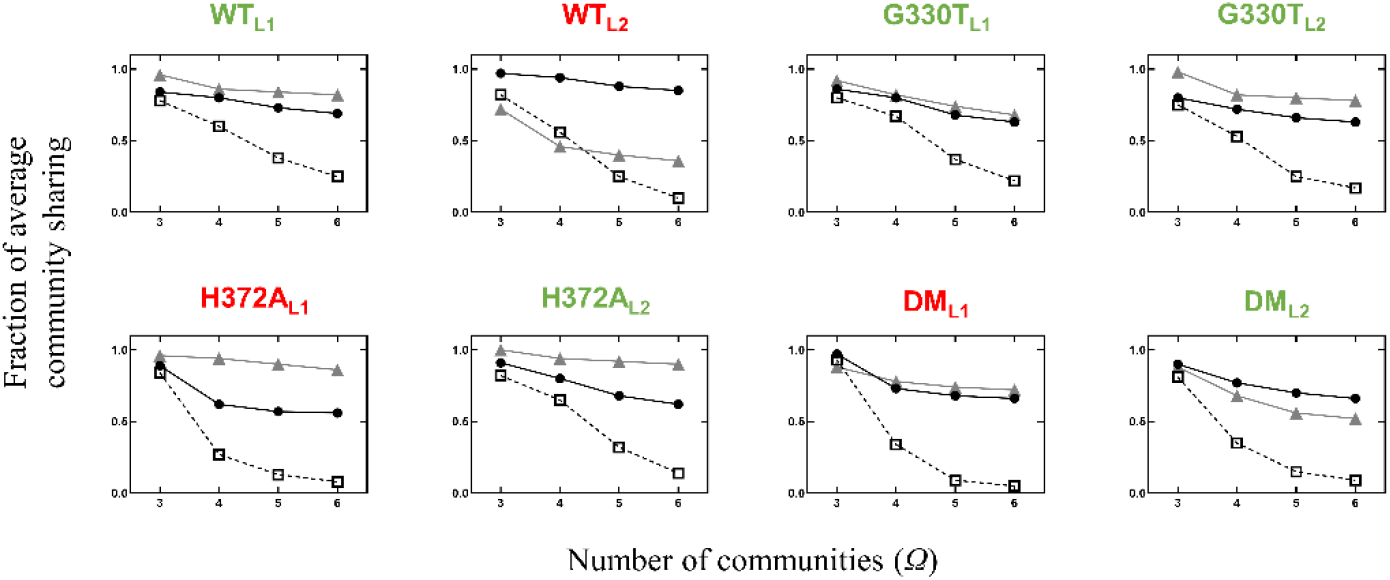
Average community co-occupancy fraction for, (◻) N-terminus, α_3_ and ligand triplet; (•) α_3_ and ligand pair in full length PDZ3 systems; (Δ) α_3_ and ligand in PDZ3^Δ^ systems.

### Major communities, dedicated membership and ubiquitous residues

Our discussion so far has utilized community sharing of structural units whereby if at least one residue from each element appears in the same community, they are listed in the co-occupancy fractions. To better explain the dynamical shifts in shared communities, we have superposed the separation into communities as a visualization on the protein structures as explained under methods. In Figure 5 we observe that all the complexes have split into three main regions. One major community organizes around the binding site, including most of the ligand and the α_2_ helix; this community is predominantly green. A second one is dominated by the α_3_ helix colored blue. A third community, colored red, includes the β_1_ strand and the surrounding loops as well as the residue at position 0 of the ligand (residue 9 in Figure 1). There are stark differences in the behavior of some other regions, however.

**Figure 5.**
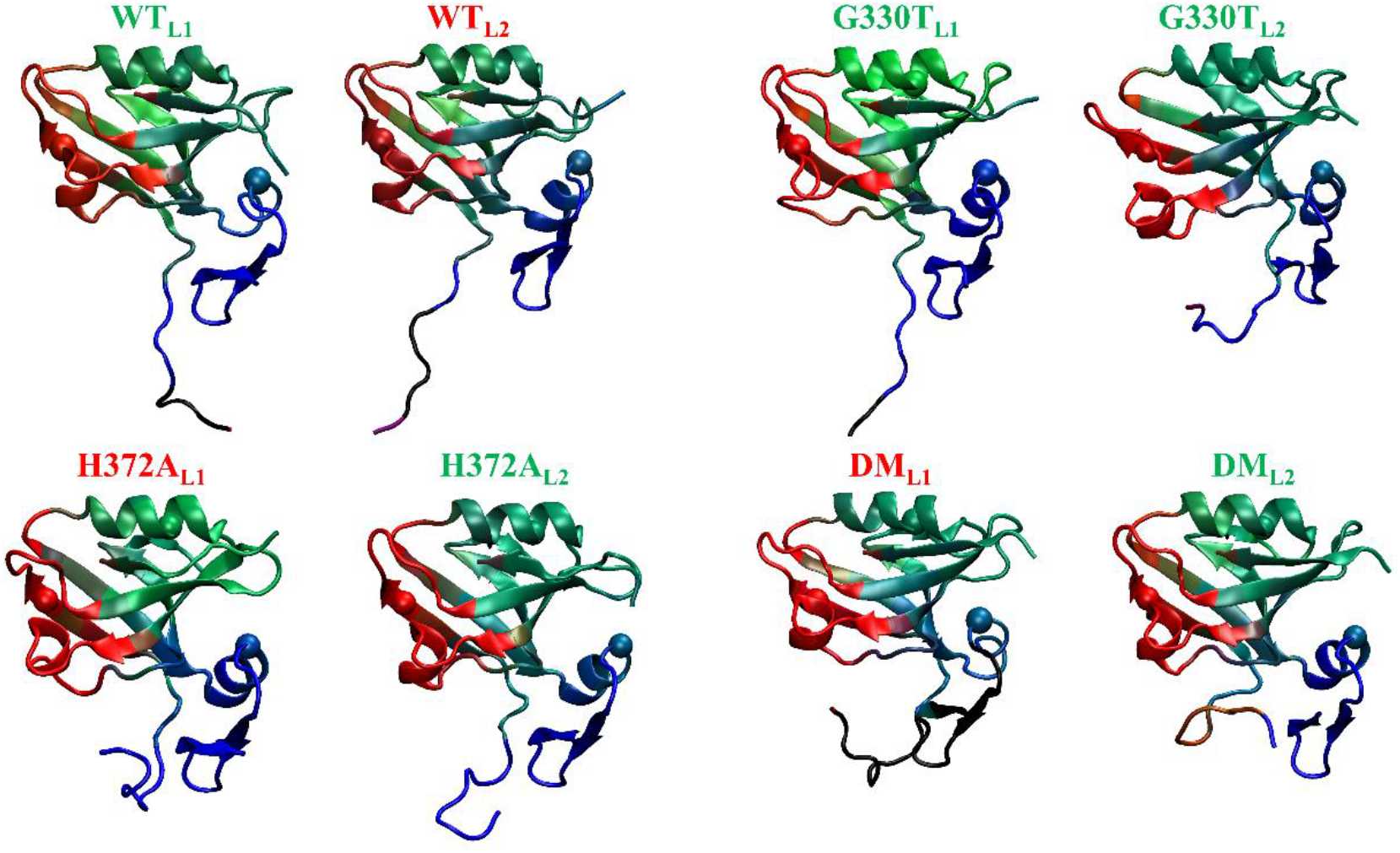
Visuals of dynamical community composition for selected variants, superimposed on the average structure for each variant. Community composition for each residue at *Ω* = 4 is accumulated throughout the trajectories (see Methods for visualization calculation details). Residues predominantly sharing the same community with I316, A375 and F400 (as ball representation) are in red, green and blue (RGB), respectively. Dynamically changing community neighbors relative to these reference residues are displayed as a mixture of RGB colors. The red community is separated out from the rest in all variants. The blue community carrying the α_3_ helix dominates the green community carrying α_2_ and the ligand since even A375 residing at the center of the latter has blue components in H372A and DM variants (hence the tinted green color). Black residues never share a community with the main selections, always separating out into a separate community.

For example, in the functional complexes WT_L1_ and G330T_L1_, the regions are well separated from each other, consisting of “pure” RGB colors except for the ligand and some parts in the central β sheet. In addition, the first few residues of the N-terminus are always separated out from the rest of the protein, indicated by their black color. In these complexes most of the N-terminus communicates with the α_3_ helix throughout the trajectory, together forming the blue region which dominates the dynamics in this variant. The shades residues take indicate they share communities with the red, green, and blue regions proportionately. In particular, except at position 0 which is red, the ligand has the shade of teal with green:blue ratio of 2, i.e., it co-inhabits the blue region a third of the time. Note that the ligand forming an ingroup with the blue region is a must for favorable complexes; e.g., in the unfavorable H372A_L1_, the ligand is pure green hence lacking a dynamical unification with the whole protein, although all other features of this complex is similar to WT_L1_ and G330T_L1_.

In G330T_L2_ and H372A_L2_, though node BC is stable throughout the MD trajectory, community compositions show that the underlying interactions change over time. In particular, the groups containing the α_2_/α_3_ helices are blended, the teal color of the former signifying 1:1 green:blue ratio, including the ligand. In this case, the whole N-terminus groups with the C-terminus and the α_3_ helix. However, the α_3_ helix itself is not a separate group but blends with the ligand/α_2_. This high grouping of the core with the N/C termini and the α_3_ helix is proposed to reinforce the high binding affinity of these variants.

The DM_L2_ complex, which is the one with the highest binding affinity to its ligand,^17^ displays a further property of community sharing. While the average number of hydrogen bonds between the ligand and PDZ3 increased from ~3.2 to 3.9 in the double mutants^47^ due to the decreased overall size of these variants, the central (green) region still communicates a great deal with the blue part.

Unique to this variant is the behavior of the N-terminus which shares its time partnering with the green and red regions reinforcing the binding by interacting with the β_1_ strand with some of its (orange colored) residues in addition to its α_3_ interactions.

Finally, it is clear from the coloring of the unfavorable WT_L2_ and DM_L1_ complexes, when the majority of the N-terminus does not co-inhabit communities with the body of the protein, favorable binding does not take place.

## CONCLUSIONS

The biological function of PDZ3 is measured by binding affinity experiments,^16, 17^ and functional complexes such as WT_L1_, G330T_L1_, G330T_L2_, H372A_L2_ and DM_L2_ are revealed; however, the underlying mechanism deciding the binding fate is not clear in this allosteric protein. The conformational differences between these forms are minimal, and a dynamical assessment is needed to further interpret the allosteric mechanism involved in communication. In a recent study on PDZ2, it was discussed that the mean structures do not necessarily differ between favorable and unfavorable complexes, whereby having different fraction of substates leads to ligand binding.^48^ Similarly, studies on PDZ3 have shown that while the conformational change is not essential for allostery,^49, 50^ N/C termini and α_3_ play an important role on its function;^5, 7–10, 20, 21^ e.g., α_3_ exhibits significantly different impact on ligand-binding within the temperature limits of 10 - 40º C.^11^ The affinity experiments demonstrate that the difference of binding free energy between the functional and dysfunctional PDZ3 complexes is around ~2.5 kcal/mol,^16^ whereby overall energy contribution of a non-bonded interaction, including hydrogen bonds, is higher than ~1.0 kcal/mol.^51–54^ Therefore, assessing conformational changes resulting from these energy differences is challenging.

In this study we develop a methodology to decipher the hidden states governing favorable binding and specificity. We set out to show that the slight variations occurring during the dynamical motions provide information on the functionality of PDZ3 complexes, and the community composition of underlying states dictates the allosteric communication without undertaking significant conformational changes.^49, 50^ For this purpose, MD-simulated trajectories of PDZ3 complexes are investigated by using community detection tools from graph theory.^35, 36^ To understand the modularity of PDZ3, the communication between the N/C termini, α_2_, α_3_ and the ligand is assessed.^7, 9, 20, 22^ Although, node centrality measures are informative to the extent of pinpointing residues whose centrality are shifted depending on the variant studied (Figure 2), our community composition analysis is more sensitive (Tables 2 and 3). We find that PDZ3 complex variants have diverse community configurations, and the fraction of these changes modulates binding preferences.

For a successful binding event, the following conditions must be met: (i) The N-terminus must share a community with the C-terminus and α_3_(blue communities in Figure 5); (ii) ligand must not only be a part of the binding site community (green communities in Figure 5), but it must also share communities with the N/C termini and α_3_ in (i). Moreover, if the N-terminus co-inhabits community with the β_1_ helix, the ligand specificity is further reinforced (orange region in Figure 5, DM_L2_). In sum, while the N-terminus confers the specificity, C-terminus and α_3_ are the essential vehicles for the formation of the PDZ complex. In fact, α_3_ acts as a hub for the whole protein by sustaining the communication with all structural segments. We propose the method developed in this work as a general methodology to study protein structures where the mechanism of action is not readily disclosed by conformational changes.

## ASSOCIATED CONTENT

### Supporting Information

Figure S1. Radial distribution function results; Figure S2. BC and MSF of each full length PDZ3 complex; Figure S3. Time evolution of number of broken edges for emergence of communities; Figure S4. Time evolution of number of community members; Table S1 Fraction of community sharing between pairs of structural segments for full length PDZ3; Table S2 Fraction of community sharing be tween pa irs of structural segme nts i n tru ncated (Δ) P DZ3.

## AUTHOR INFORMATION

## Author Contributions

T.F.G. designed the research, conducted the computer experiments, analyzed results, interpreted data, wrote the paper, and constructed figures. A.R.A. and C.A. designed the research, guided the computer experiments and analyses, interpreted data, guided the structure and contents of the paper, and edited the paper. All authors have given approval to the final version of the manuscript.

## Supporting Information

**Table S1.** Fraction of instances pairs of structural segments co-inhabit a community.*

| System label | N-term and C-term | | | | N-term and $\alpha_3$ | | | | N-term and $\alpha_2$ | | | |
| --- | --- | --- | --- | --- | --- | --- | --- | --- | --- | --- | --- | --- |
| | $\Omega = 3$ | $\Omega = 4$ | $\Omega = 5$ | $\Omega = 6$ | $\Omega = 3$ | $\Omega = 4$ | $\Omega = 5$ | $\Omega = 6$ | $\Omega = 3$ | $\Omega = 4$ | $\Omega = 5$ | $\Omega = 6$ |
| **WT<sub>L1</sub>** | **1.00 ± 0.01** | **1.00 ± 0.01** | **0.98 ± 0.01** | **0.97 ± 0.01** | **0.99 ± 0.01** | **0.96 ± 0.01** | **0.88 ± 0.02** | **0.78 ± 0.03** | **0.85 ± 0.02** | 0.60 ± 0.03 | **0.33 ± 0.03** | **0.16 ± 0.02** |
| **WT<sub>L2</sub>** | **0.80 ± 0.03** | **0.79 ± 0.03** | **0.75 ± 0.03** | **0.70 ± 0.03** | **0.89 ± 0.02** | **0.84 ± 0.02** | **0.73 ± 0.03** | 0.67 ± 0.03 | **0.81 ± 0.03** | 0.61 ± 0.03 | **0.33 ± 0.03** | **0.12 ± 0.02** |
| **G330T<sub>L1</sub>** | **0.97 ± 0.01** | **0.97 ± 0.01** | **0.96 ± 0.01** | **0.95 ± 0.01** | **0.95 ± 0.01** | **0.90 ± 0.02** | **0.85 ± 0.02** | **0.83 ± 0.02** | **0.87 ± 0.02** | 0.68 ± 0.03 | **0.31 ± 0.03** | **0.11 ± 0.02** |
| **G330T<sub>L2</sub>** | **0.92 ± 0.02** | **0.91 ± 0.02** | **0.88 ± 0.02** | **0.85 ± 0.02** | **0.97 ± 0.01** | **0.93 ± 0.02** | **0.86 ± 0.02** | **0.79 ± 0.03** | **0.86 ± 0.02** | 0.62 ± 0.03 | **0.35 ± 0.03** | **0.17 ± 0.02** |
| **H372A<sub>L1</sub>** | **1.00 ± 0.01** | **0.99 ± 0.01** | **0.99 ± 0.01** | **0.98 ± 0.01** | **0.98 ± 0.01** | **0.96 ± 0.01** | **0.91 ± 0.02** | **0.83 ± 0.02** | **0.85 ± 0.02** | **0.43 ± 0.03** | **0.28 ± 0.03** | **0.15 ± 0.02** |
| **H372A<sub>L2</sub>** | **1.00 ± 0.01** | **1.00 ± 0.01** | **1.00 ± 0.01** | **1.00 ± 0.01** | **0.96 ± 0.01** | **0.95 ± 0.01** | **0.94 ± 0.02** | **0.90 ± 0.02** | **0.86 ± 0.02** | 0.68 ± 0.03 | **0.38 ± 0.03** | **0.17 ± 0.02** |
| **DM<sub>L1</sub>** | **1.00 ± 0.01** | **1.00 ± 0.01** | **0.99 ± 0.01** | **0.99 ± 0.01** | **0.99 ± 0.01** | **0.94 ± 0.02** | **0.82 ± 0.02** | **0.72 ± 0.03** | **0.92 ± 0.02** | **0.46 ± 0.03** | **0.19 ± 0.03** | **0.10 ± 0.02** |
| **DM<sub>L2</sub>** | **0.99 ± 0.01** | **0.98 ± 0.01** | **0.97 ± 0.01** | **0.96 ± 0.01** | **0.99 ± 0.01** | **0.98 ± 0.01** | **0.95 ± 0.01** | **0.94 ± 0.02** | **0.86 ± 0.02** | **0.41 ± 0.03** | **0.21 ± 0.03** | **0.13 ± 0.02** |
| System label | C-term and ligand | | | | C-term and $\alpha_3$ | | | | C-term and $\alpha_2$ | | | |
| | $\Omega = 3$ | $\Omega = 4$ | $\Omega = 5$ | $\Omega = 6$ | $\Omega = 3$ | $\Omega = 4$ | $\Omega = 5$ | $\Omega = 6$ | $\Omega = 3$ | $\Omega = 4$ | $\Omega = 5$ | $\Omega = 6$ |
| **WT<sub>L1</sub>** | **0.38 ± 0.03** | **0.35 ± 0.03** | **0.32 ± 0.03** | **0.31 ± 0.03** | **0.95 ± 0.01** | **0.95 ± 0.01** | **0.95 ± 0.01** | **0.95 ± 0.01** | **0.17 ± 0.02** | **0.07 ± 0.02** | **0.03 ± 0.01** | **0.01 ± 0.01** |
| **WT<sub>L2</sub>** | **0.35 ± 0.03** | **0.31 ± 0.03** | **0.28 ± 0.03** | **0.26 ± 0.03** | **0.96 ± 0.01** | **0.95 ± 0.01** | **0.95 ± 0.01** | **0.95 ± 0.01** | **0.28 ± 0.03** | **0.17 ± 0.02** | **0.12 ± 0.02** | **0.11 ± 0.02** |
| **G330T<sub>L1</sub>** | **0.42 ± 0.03** | **0.39 ± 0.03** | **0.37 ± 0.03** | **0.35 ± 0.03** | **0.97 ± 0.01** | **0.96 ± 0.01** | **0.96 ± 0.01** | **0.95 ± 0.01** | **0.18 ± 0.03** | **0.07 ± 0.02** | **0.03 ± 0.01** | **0.01 ± 0.01** |
| **G330T<sub>L2</sub>** | **0.31 ± 0.03** | **0.24 ± 0.03** | **0.23 ± 0.03** | **0.21 ± 0.03** | **0.96 ± 0.01** | **0.96 ± 0.01** | **0.96 ± 0.01** | **0.96 ± 0.01** | **0.26 ± 0.03** | **0.10 ± 0.02** | **0.06 ± 0.02** | **0.04 ± 0.01** |
| **H372A<sub>L1</sub>** | **0.26 ± 0.03** | **0.15 ± 0.02** | **0.11 ± 0.02** | **0.09 ± 0.02** | **0.92 ± 0.02** | **0.92 ± 0.02** | **0.91 ± 0.02** | **0.91 ± 0.02** | **0.22 ± 0.03** | **0.10 ± 0.02** | **0.04 ± 0.01** | **0.01 ± 0.01** |
| **H372A<sub>L2</sub>** | **0.27 ± 0.03** | **0.22 ± 0.03** | **0.19 ± 0.03** | **0.17 ± 0.02** | **0.98 ± 0.01** | **0.98 ± 0.01** | **0.98 ± 0.01** | **0.98 ± 0.01** | **0.15 ± 0.02** | **0.08 ± 0.02** | **0.04 ± 0.01** | **0.02 ± 0.01** |
| **DM<sub>L1</sub>** | **0.48 ± 0.03** | **0.25 ± 0.03** | **0.17 ± 0.02** | **0.15 ± 0.02** | **0.91 ± 0.02** | **0.91 ± 0.02** | **0.91 ± 0.02** | **0.91 ± 0.02** | **0.46 ± 0.03** | **0.20 ± 0.03** | **0.10 ± 0.02** | **0.08 ± 0.02** |
| **DM<sub>L2</sub>** | **0.23 ± 0.03** | **0.17 ± 0.02** | **0.16 ± 0.02** | **0.14 ± 0.02** | **0.98 ± 0.01** | **0.98 ± 0.01** | **0.98 ± 0.01** | **0.98 ± 0.01** | **0.19 ± 0.03** | **0.10 ± 0.02** | **0.07 ± 0.02** | **0.05 ± 0.01** |
| System label | $\alpha_3$ and ligand | | | | $\alpha_3$ and $\alpha_2$ | | | | $\alpha_2$ and ligand | | | |
| | $\Omega = 3$ | $\Omega = 4$ | $\Omega = 5$ | $\Omega = 6$ | $\Omega = 3$ | $\Omega = 4$ | $\Omega = 5$ | $\Omega = 6$ | $\Omega = 3$ | $\Omega = 4$ | $\Omega = 5$ | $\Omega = 6$ |
| **WT<sub>L1</sub>** | **0.84 ± 0.02** | **0.80 ± 0.03** | **0.73 ± 0.03** | 0.69 ± 0.03 | **0.74 ± 0.03** | 0.58 ± 0.03 | **0.41 ± 0.03** | **0.28 ± 0.03** | **0.99 ± 0.01** | **0.92 ± 0.02** | **0.86 ± 0.02** | **0.80 ± 0.03** |
| **WT<sub>L2</sub>** | **0.97 ± 0.01** | **0.94 ± 0.02** | **0.88 ± 0.02** | **0.85 ± 0.02** | **0.96 ± 0.01** | **0.83 ± 0.02** | **0.76 ± 0.03** | 0.64 ± 0.03 | **1.00 ± 0.01** | **0.95 ± 0.01** | **0.94 ± 0.02** | **0.89 ± 0.02** |
| **G330T<sub>L1</sub>** | **0.86 ± 0.02** | **0.80 ± 0.03** | 0.68 ± 0.03 | 0.63 ± 0.03 | **0.77 ± 0.03** | 0.55 ± 0.03 | **0.35 ± 0.03** | **0.23 ± 0.03** | **0.99 ± 0.01** | **0.93 ± 0.02** | **0.85 ± 0.02** | **0.78 ± 0.03** |
| **G330T<sub>L2</sub>** | **0.80 ± 0.03** | **0.72 ± 0.03** | 0.66 ± 0.03 | 0.63 ± 0.03 | **0.75 ± 0.03** | 0.54 ± 0.03 | **0.44 ± 0.03** | **0.34 ± 0.03** | **0.98 ± 0.01** | **0.91 ± 0.02** | **0.86 ± 0.02** | **0.82 ± 0.03** |
| **H372A<sub>L1</sub>** | **0.89 ± 0.02** | 0.62 ± 0.03 | 0.57 ± 0.03 | 0.56 ± 0.03 | **0.87 ± 0.02** | 0.65 ± 0.03 | 0.58 ± 0.03 | 0.51 ± 0.03 | **1.00 ± 0.01** | **1.00 ± 0.01** | **1.00 ± 0.01** | **1.00 ± 0.01** |
| **H372A<sub>L2</sub>** | **0.91 ± 0.02** | **0.80 ± 0.03** | 0.68 ± 0.03 | 0.62 ± 0.03 | **0.88 ± 0.02** | **0.75 ± 0.03** | 0.64 ± 0.03 | 0.54 ± 0.03 | **1.00 ± 0.01** | **1.00 ± 0.01** | **0.98 ± 0.01** | **0.96 ± 0.01** |
| **DM<sub>L1</sub>** | **0.97 ± 0.01** | **0.73 ± 0.03** | 0.68 ± 0.03 | 0.66 ± 0.03 | **0.95 ± 0.01** | 0.67 ± 0.03 | 0.58 ± 0.03 | 0.54 ± 0.03 | **1.00 ± 0.01** | **1.00 ± 0.01** | **0.99 ± 0.01** | **0.98 ± 0.01** |
| **DM<sub>L2</sub>** | **0.90 ± 0.02** | **0.77 ± 0.03** | **0.70 ± 0.03** | 0.66 ± 0.03 | **0.85 ± 0.02** | 0.67 ± 0.03 | 0.57 ± 0.03 | 0.50 ± 0.03 | **1.00 ± 0.01** | **0.99 ± 0.01** | **0.98 ± 0.01** | **0.97 ± 0.01** |

\*values greater than 0.7 shown in bold; those less than 0.5 are colored blue. N-terminus–ligand values are reported in Table 3 of the main text.

**Table S2.** Fraction of instances pairs of structural segments co-inhabit a community in truncated (Δ) PDZ3.*

| System label | C-term and ligand | | | | C-term and $\alpha_3$ | | | | C-term and $\alpha_2$ | | | |
| --- | --- | --- | --- | --- | --- | --- | --- | --- | --- | --- | --- | --- |
| | $\Omega = 3$ | $\Omega = 4$ | $\Omega = 5$ | $\Omega = 6$ | $\Omega = 3$ | $\Omega = 4$ | $\Omega = 5$ | $\Omega = 6$ | $\Omega = 3$ | $\Omega = 4$ | $\Omega = 5$ | $\Omega = 6$ |
| **WT**<sub>L1</sub><sup>Δ</sup> | 0.20 ± 0.06 | 0.18 ± 0.05 | 0.18 ± 0.05 | 0.16 ± 0.05 | **0.98 ± 0.02** | **0.98 ± 0.02** | **0.98 ± 0.02** | **0.98 ± 0.02** | **0.08 ± 0.03** | 0.06 ± 0.03 | 0.04 ± 0.03 | 0.02 ± 0.02 |
| **WT**<sub>L2</sub><sup>Δ</sup> | **0.02 ± 0.02** | **0.02 ± 0.02** | **0.02 ± 0.02** | **0.02 ± 0.02** | **1.00 ± 0.01** | **1.00 ± 0.01** | **1.00 ± 0.01** | **1.00 ± 0.01** | 0 | 0 | 0 | 0 |
| **G330T**<sub>L1</sub><sup>Δ</sup> | **0.22 ± 0.06** | **0.22 ± 0.06** | **0.20 ± 0.06** | **0.18 ± 0.05** | **1.00 ± 0.01** | **1.00 ± 0.01** | **1.00 ± 0.01** | **1.00 ± 0.01** | 0.16 ± 0.05 | 0.06 ± 0.03 | 0.06 ± 0.03 | 0.04 ± 0.03 |
| **G330T**<sub>L2</sub><sup>Δ</sup> | 0.26 ± 0.06 | 0.22 ± 0.06 | 0.22 ± 0.06 | 0.16 ± 0.05 | **1.00 ± 0.01** | **1.00 ± 0.01** | **1.00 ± 0.01** | **1.00 ± 0.01** | **0.02 ± 0.02** | 0 | 0 | 0 |
| **H372A**<sub>L1</sub><sup>Δ</sup> | 0.16 ± 0.05 | 0.16 ± 0.05 | 0.10 ± 0.04 | 0.06 ± 0.03 | **0.98 ± 0.02** | **0.98 ± 0.02** | **0.98 ± 0.02** | **0.98 ± 0.02** | **0.06 ± 0.03** | **0.04 ± 0.03** | 0.02 ± 0.02 | 0.02 ± 0.02 |
| **H372A**<sub>L2</sub><sup>Δ</sup> | **0.54 ± 0.07** | **0.52 ± 0.07** | **0.38 ± 0.07** | **0.36 ± 0.07** | **0.98 ± 0.02** | **0.98 ± 0.02** | **0.98 ± 0.02** | **0.98 ± 0.02** | **0.32 ± 0.07** | **0.26 ± 0.06** | **0.22 ± 0.06** | **0.18 ± 0.05** |
| **DM**<sub>L1</sub><sup>Δ</sup> | **0.16 ± 0.05** | **0.16 ± 0.05** | 0.14 ± 0.05 | 0.10 ± 0.04 | **0.96 ± 0.03** | **0.96 ± 0.03** | **0.96 ± 0.03** | **0.96 ± 0.03** | **0.10 ± 0.04** | **0.08 ± 0.04** | 0.06 ± 0.03 | **0.02 ± 0.02** |
| **DM**<sub>L2</sub><sup>Δ</sup> | 0.16 ± 0.05 | 0.14 ± 0.05 | 0.08 ± 0.04 | 0.06 ± 0.03 | **0.96 ± 0.03** | **0.96 ± 0.03** | **0.96 ± 0.03** | **0.96 ± 0.03** | 0.14 ± 0.05 | 0.10 ± 0.02 | 0.02 ± 0.02 | 0.02 ± 0.02 |
| System label | $\alpha_3$ and ligand | | | | $\alpha_3$ and $\alpha_2$ | | | | $\alpha_2$ and ligand | | | |
| | $\Omega = 3$ | $\Omega = 4$ | $\Omega = 5$ | $\Omega = 6$ | $\Omega = 3$ | $\Omega = 4$ | $\Omega = 5$ | $\Omega = 6$ | $\Omega = 3$ | $\Omega = 4$ | $\Omega = 5$ | $\Omega = 6$ |
| **WT**<sub>L1</sub><sup>Δ</sup> | **0.96 ± 0.03** | **0.86 ± 0.05** | **0.84 ± 0.05** | **0.82 ± 0.05** | **0.82 ± 0.05** | 0.62 ± 0.07 | 0.44 ± 0.07 | 0.24 ± 0.06 | **0.92 ± 0.04** | **0.76 ± 0.06** | 0.68 ± 0.07 | 0.60 ± 0.07 |
| **WT**<sub>L2</sub><sup>Δ</sup> | **0.72 ± 0.06** | 0.46 ± 0.07 | 0.40 ± 0.07 | 0.36 ± 0.07 | 0.66 ± 0.07 | 0.44 ± 0.07 | 0.30 ± 0.07 | 0.22 ± 0.06 | **0.98 ± 0.02** | **0.94 ± 0.03** | **0.90 ± 0.04** | **0.86 ± 0.05** |
| **G330T**<sub>L1</sub><sup>Δ</sup> | **0.92 ± 0.04** | **0.82 ± 0.05** | 0.74 ± 0.06 | 0.68 ± 0.07 | **0.86 ± 0.05** | 0.46 ± 0.07 | 0.18 ± 0.05 | 0.12 ± 0.05 | **1.00 ± 0.01** | **0.80 ± 0.06** | **0.70 ± 0.07** | 0.66 ± 0.07 |
| **G330T**<sub>L2</sub><sup>Δ</sup> | **0.98 ± 0.02** | **0.82 ± 0.05** | **0.80 ± 0.06** | 0.78 ± 0.06 | **0.90 ± 0.04** | **0.72 ± 0.06** | 0.52 ± 0.07 | 0.38 ± 0.07 | **0.98 ± 0.02** | **0.94 ± 0.03** | **0.92 ± 0.04** | **0.92 ± 0.04** |
| **H372A**<sub>L1</sub><sup>Δ</sup> | **0.96 ± 0.03** | **0.94 ± 0.03** | **0.90 ± 0.04** | **0.86 ± 0.05** | **0.90 ± 0.04** | **0.80 ± 0.06** | 0.64 ± 0.07 | 0.56 ± 0.07 | **1.00 ± 0.01** | **0.92 ± 0.04** | **0.90 ± 0.04** | **0.88 ± 0.05** |
| **H372A**<sub>L2</sub><sup>Δ</sup> | **1.00 ± 0.01** | **0.94 ± 0.03** | **0.92 ± 0.04** | **0.90 ± 0.04** | **0.96 ± 0.03** | **0.86 ± 0.05** | **0.78 ± 0.06** | 0.66 ± 0.07 | **1.00 ± 0.01** | **0.94 ± 0.03** | **0.90 ± 0.04** | **0.86 ± 0.05** |
| **DM**<sub>L1</sub><sup>Δ</sup> | **0.88 ± 0.05** | **0.78 ± 0.06** | **0.74 ± 0.06** | **0.72 ± 0.06** | **0.84 ± 0.05** | **0.80 ± 0.06** | 0.64 ± 0.07 | 0.50 ± 0.07 | **1.00 ± 0.01** | **0.98 ± 0.02** | **0.96 ± 0.03** | **0.96 ± 0.03** |
| **DM**<sub>L2</sub><sup>Δ</sup> | **0.88 ± 0.05** | 0.68 ± 0.07 | **0.56 ± 0.07** | **0.52 ± 0.07** | **0.84 ± 0.05** | 0.68 ± 0.07 | **0.46 ± 0.07** | **0.38 ± 0.07** | **1.00 ± 0.01** | **1.00 ± 0.01** | **1.00 ± 0.01** | **0.98 ± 0.02** |

\*values greater than 0.7 shown in bold; those differing from the full-length variant value (Table S1) by more than the sum of the error bars colored red.

**Figure S1.**
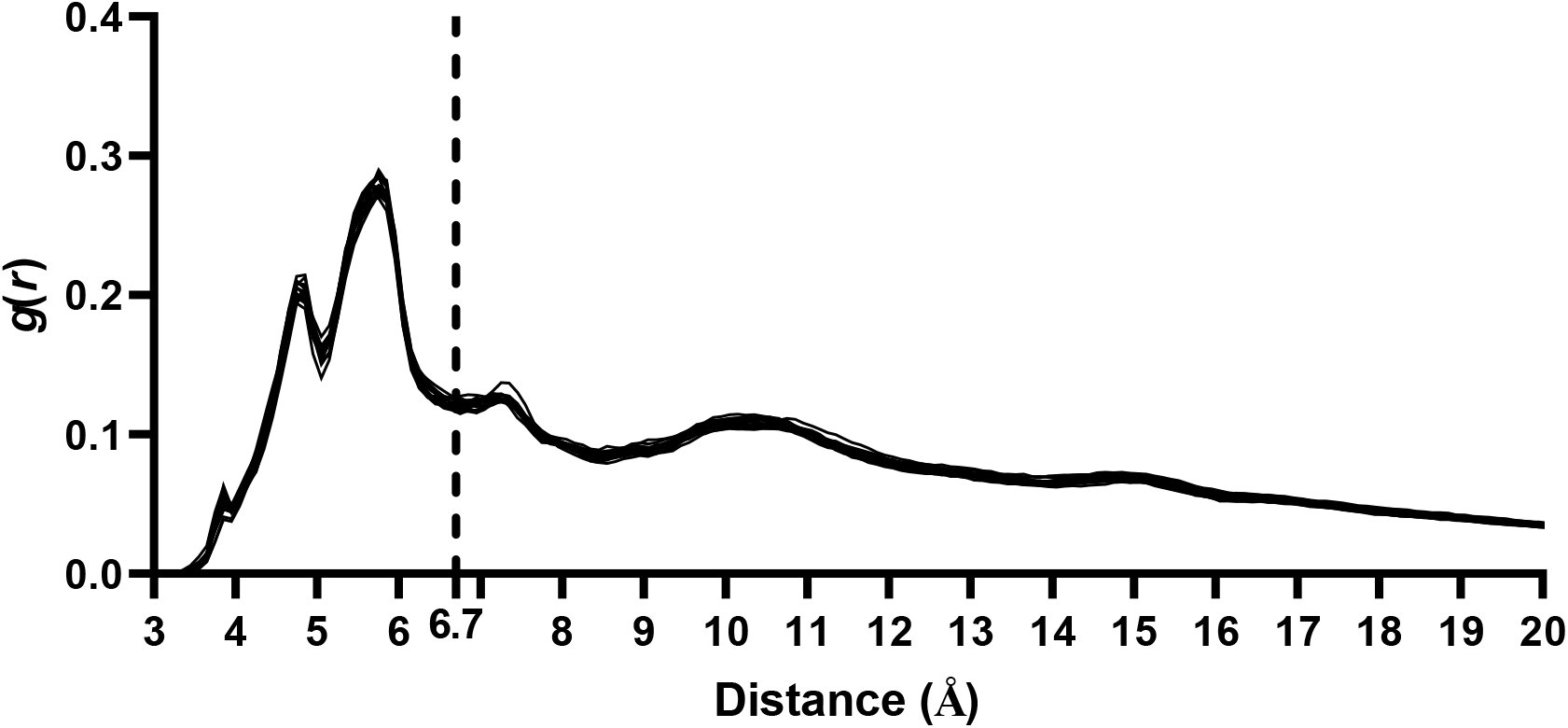
Radial distribution function, *g*(*r*), between residue centers, results from the 16 simulations belonging to all full length PDZ3 complexes are superimposed. The cut-off distance of 6.7 Å signified by dashed line marks the completion location of the first coordination shell of non-bonded C_β_ atoms; this cut-off is utilized for all network analyses in this study. Note that selection of C_β_ atoms as coarse-graining centers is essential to capture dynamical communication between residue side chains.

**Figure S2.**
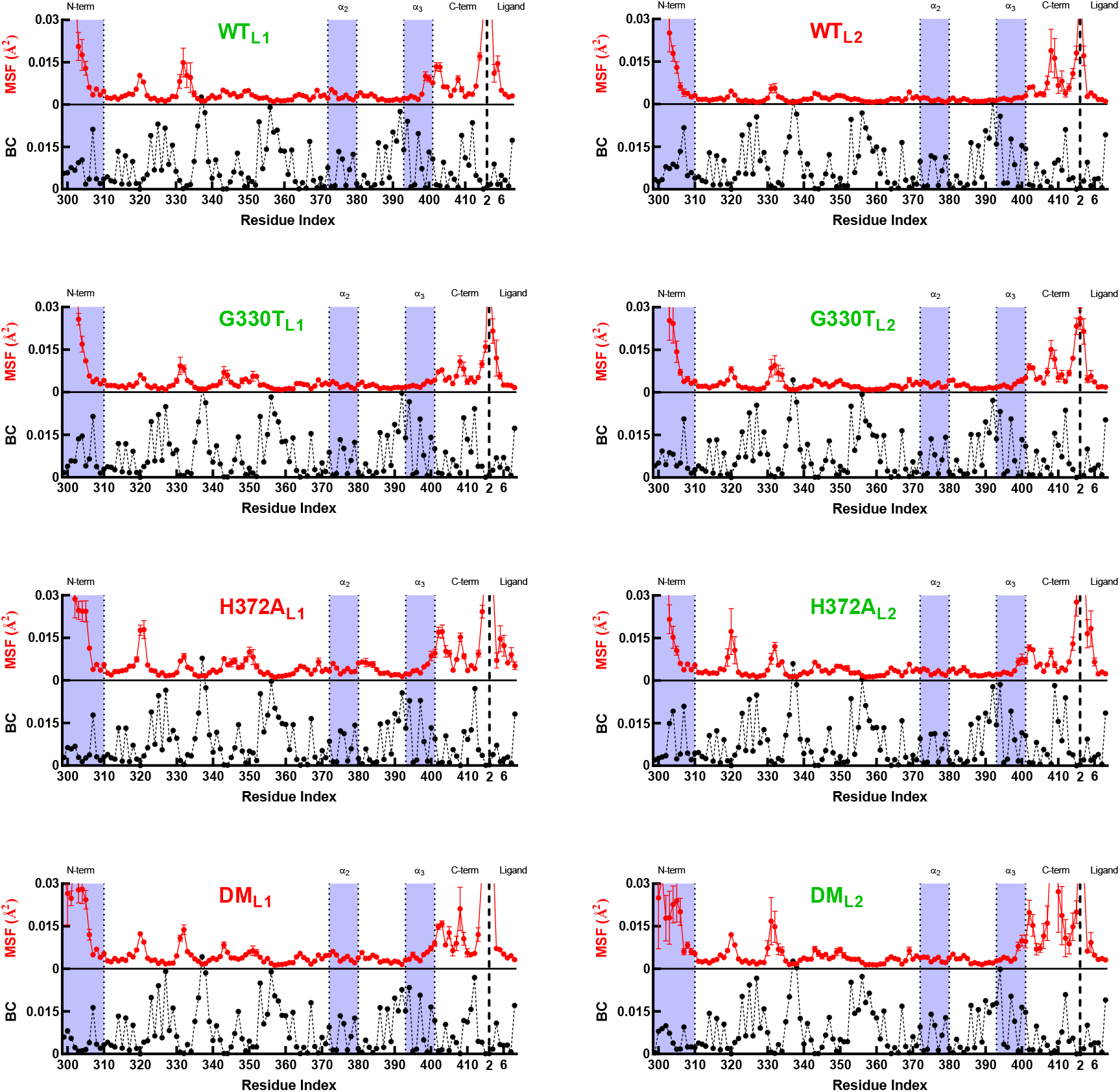
BC and MSF of each PDZ3 complex and ΔBC through single mutation paths. Structural segments are labeled at the top of the graph, and segments of interest are highlighted. Favorable and unfavorable complexes are labelled by green and red, respectively. MSF results are computed by using C_β_ atoms of 40-ns long chunks of MD trajectories in the 120-200 ns time window; thus, six chunks are averaged and error bars are displayed.

**Figure S3.**
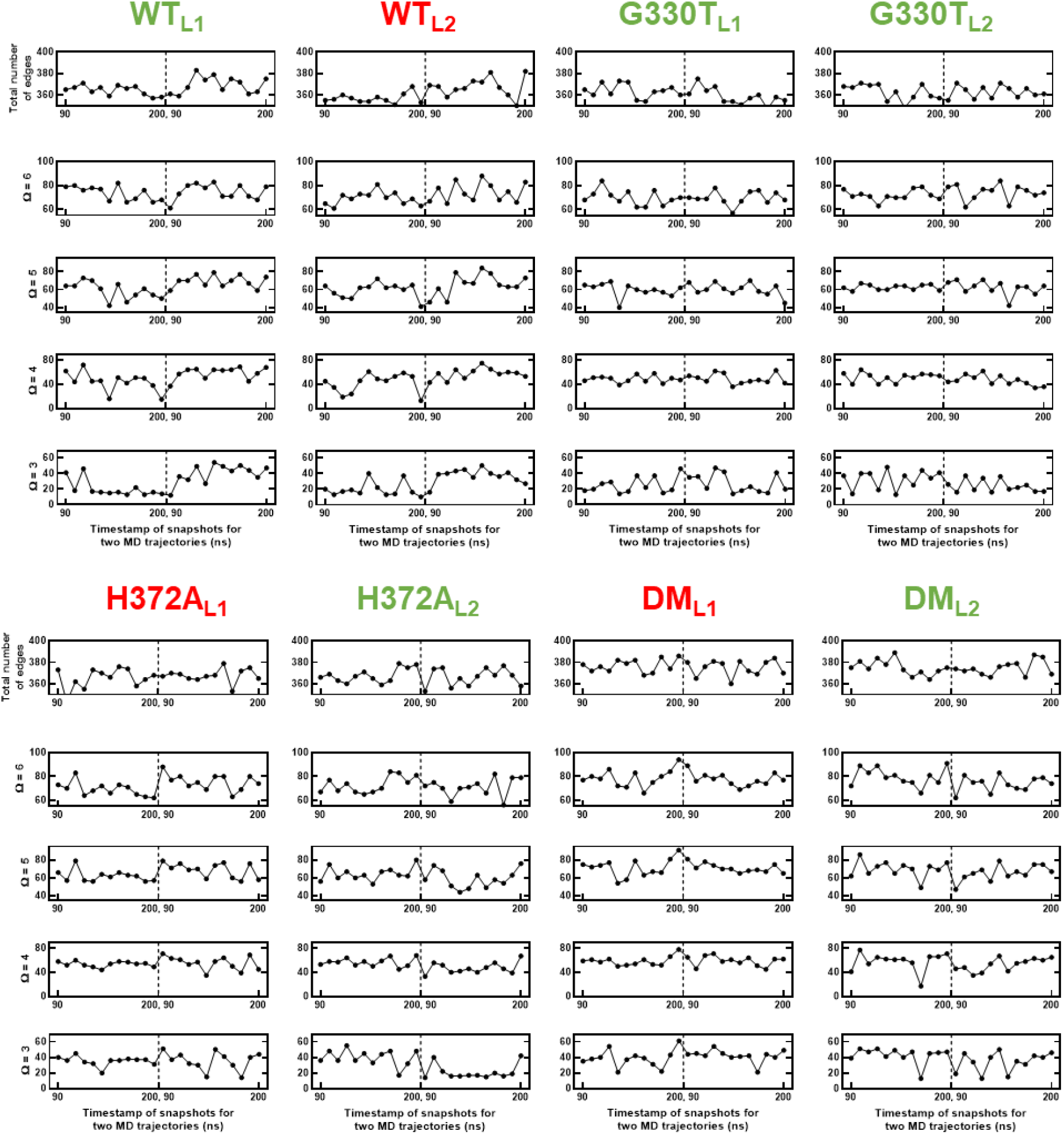
Time evolution of the number of broken edges for emergence of *Ω* =3-6 communities. Grand averages are reported in Table 2.

**Figure S4.**
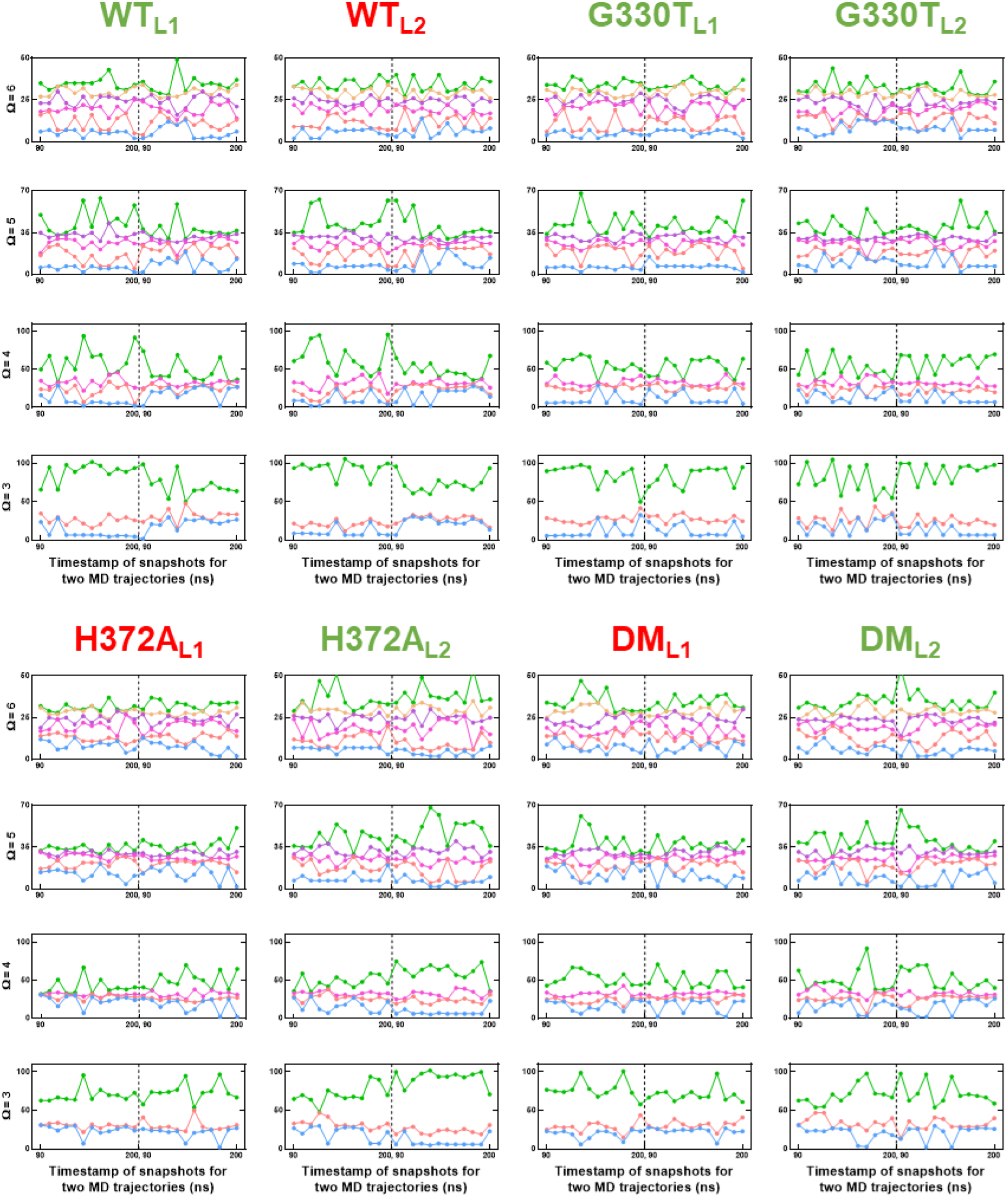
Time evolution of number of community members for *Ω* =3-6 communities. Smallest community is shown in blue, while the largest in green.

